# Inferring Metabolic States from Single Cell Transcriptomic Data via Geometric Deep Learning

**DOI:** 10.1101/2023.12.05.570153

**Authors:** Holly Steach, Siddharth Viswanath, Yixuan He, Xitong Zhang, Natalia Ivanova, Matthew Hirn, Michael Perlmutter, Smita Krishnaswamy

## Abstract

The ability to measure gene expression at single-cell resolution has elevated our understanding of how biological features emerge from complex and interdependent networks at molecular, cellular, and tissue scales. As technologies have evolved that complement scRNAseq measurements with things like single-cell proteomic, epigenomic, and genomic information, it becomes increasingly apparent how much biology exists as a product of multimodal regulation. Biological processes such as transcription, translation, and post-translational or epigenetic modification impose both energetic and specific molecular demands on a cell and are therefore implicitly constrained by the metabolic state of the cell. While metabolomics is crucial for defining a holistic model of any biological process, the chemical heterogeneity of the metabolome makes it particularly difficult to measure, and technologies capable of doing this at single-cell resolution are far behind other multiomics modalities. To address these challenges, we present GEFMAP (Gene Expression-based Flux Mapping and Metabolic Pathway Prediction), a method based on geometric deep learning for predicting flux through reactions in a global metabolic network using transcriptomics data, which we ultimately apply to scRNAseq. GEFMAP leverages the natural graph structure of metabolic networks to learn both a biological objective for each cell and estimate a mass-balanced relative flux rate for each reaction in each cell using novel deep learning models.

## 1 Introduction

Technologies that measure gene expression at single-cell resolution have massively expanded over the last decade and are now broadly available to researchers. However, at this time, no method exists that accurately measures the entire metabolome at single-cell resolution. One possible remedy is to utilize the wide availability of single-cell RNA sequencing data (scRNAseq) to develop computational tools for predicting and modeling the metabolic states of individual cells via the transcriptome. Indeed, metabolic reactions are catalyzed by enzymes whose expression is available in the scRNA-seq measurements. Here, we use this idea to understand not only what gene pathways are being upregulated by the cell but also gain specific insights on the metabolic pathways being actively regulated as a result of the gene expression. Additionally, after formulating a metabolic objective, we can compute flux through the entire network and assess the effects of transcription network-wide.

Learning about the metabolome is particularly challenging for multicellular organisms in part because we do not know a priori what objective each cell is trying to optimize. This is in contrast to the setting of single-cell organisms where researchers have effectively used cell growth (biomass accumulation) as an objective [22]. This motivates us to introduce GEFMAP - Gene Expression-based Flux Mapping and Metabolic Pathway Prediction. GEFMAP consists of two sub-networks, the first of which infers a plausible metabolic objective from the upregulated portions of the transcriptomic profile, and the second of which solves this objective in order to infer network-wide metabolic flux rates.

The first subnetwork *infers the cellular metabolic objective* based on the intuition that the cell upregulates expression of catalytic enzymes (genes) for producing its desired metabolic state. Here, we formulate this as the problem of finding a highly-weighted, highly-connected subgraph in the metabolic network graph where the nodes representing individual reactions are given weights according to the expression levels of associated genes. This allows us to essentially infer the cellular objective from its transcriptomic profile. To do this, we utilize a deep neural network based on the geometric scattering transform [5,6,32] to estimate a large highly-connected subnetwork by solving a relaxed version of the maximum weighted clique problem. We then formulate a cellular objective function corresponding to maximizing the reactivity in this subgraph.

Our second subnetwork *solves the cellular objective* given the constraint that our solution **v** must lie in the null space of the metabolic reaction stoichiometric matrix *S*. Therefore, we consider a basis for this null space, *S***v** = 0, and design a novel network that operates in this null space to find the coefficients of the solution with respect to this basis. Thus, GEFMAP is able to utilize both the structure of the network and the geometric constraint that **v** lies within the null space of *S* to predict the metabolic fluxes. In essence this allows us to predict the entire metabolic state based on the inferred objective, which will include maximizing reactions of a subnetwork and may have pervasive effects of system-wide flux.

**Main Contributions.** The main contributions in this paper are summarized as follows:

1. We create a dual system of neural networks to both infer a cellular metabolic objective as well as predict fluxes that result from the metabolic objective using deep neural network models.
2. For inferring a transcriptionally upregulated cellular objective, we modify a graph neural network designed to compute a maximum clique, into a maximum weighted subgraph.
3. For solving the objective, we use a neural network, which is constrained to preserve mass balance, to estimate the reaction rates across the network that collectively maximize flux through the set of reactions corresponding to the objective.
4. We apply these networks to various data sets to show that we are able to determine the cell’s metabolic objective and estimate flux rate in a synthetic and augmented E. coli data set.

## 2 Problem Setup

The difficulty of direct metabolomics measurements, particularly at single-cell resolution, motivates the development of computational methods that leverage widely available transcriptomic data to estimate metabolic network states. A large portion of metabolic reactions are catalyzed by enzymes, where expression of reaction-associated genes is a rate-limiting factor. A given cell will therefore transcriptionally upregulate metabolic genes in order to increase activity of a particular set of reactions in response to some stimulus. However, the relationship between transcript levels and reaction rates is influenced by several other features and is not a sufficient predictor of flux. Therefore, in order to model cellular metabolic activity from transcriptional data we define two major challenges. The first is to identify a set of reactions that a cell is actively engaging, which we refer to as the **cellular metabolomic objective**. Once known, the second challenge is to determine how the cell is accomplishing this objective, which we do by predicting flux rates for all reactions in the metabolic network. By subjecting these predictions to a set of constraints that include expression of reaction-associated genes and mass conservation within the system, we identify which reactions support the cellular objective and are likely to have high activity and which reactions are counter to the objective and are likely to be competitively downregulated.

Since metabolites and their fluxes form a network, all components of this problem can be modeled using graph representations. Here, we leverage previously curated genome-scale metabolic models (GSMM) that represent the global set of reactions, the substrate/product stoichiometries, and the associated genes involved. Tools such as constraint-based reconstruction and analysis (COBRA) [8] include methods to parse GSMM, however, we require a non-standard method of mapping gene expression to reactions. A single reaction may be catalyzed by a single gene; however, many reactions involve multiple enzymes. In the latter setting, it is sometimes the case that all of the enzymes are necessary, such as in the case of an enzyme complex, and other times it is the case that the enzymes act redundantly. Therefore, the models must be parsed in a way that we can use to map gene expression values onto reactions in a manner that reflects their biological relationships. To do this, we build a method that extracts rules from a given GSMM and applies them to expression value assignments.

A common method for predicting metabolic network activity is flux balance analysis (FBA)[22], in which flux through a metabolic network is formulated as a linear optimization problem. In this framework, we consider a system of *m* metabolites 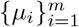 and *n* reactions 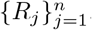, and we let *S* denote and *m × n* stoichiometry matrix so that *S*_*i*,*j*_ is the activity level of the metabolite *μ*_*i*_ in the reaction *R*_*j*_. Generally *S* is provided as part of a curated genome-scale metabolic model (GEM). We let **v** denote an *n ×* 1 flux vector where *v*_*j*_ denotes the rate of the reaction *R*_*j*_ (metabolite concentration per unit of time). Our goal is to estimate **v** based on *S* and other available information. In FBA, we assume that **v** is the solution to an optimization problem of the form, 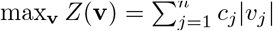, subject to certain constraints.

By construction,

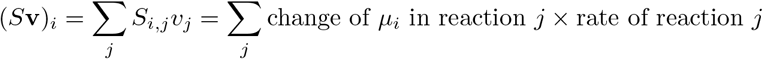

is the overall net rate change of *μ*_*i*_ in the system. In our analysis, we will assume that the system is at steady state, i.e., the mass entering the system (cellular import) is equal to the mass accumulating in or leaving the system (growth, storage, cellular export). Thus, **v** must satisfy the constraint

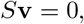

i.e., **v** must lie in the the null space of *S*. Additionally, we often have upper and lower constraints for each of the rates *v*_*j*_ based on known biological features such as environmental nutrient availability or gene expression, i.e.,

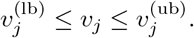

Notably, in the case where *R*_*j*_ is reversible, we will take 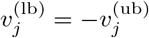 and otherwise we will have 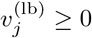.

Various cell types in different states such as quiescence, activation, and differentiation engage different metabolic pathways to support their respective energetic and biomolecular needs. These metabolic programs comprise subsets of the global metabolic network and can be modeled as an objective function that the cell is optimizing under constraints parameterized by the external environment and bioenergetic features such as reaction kinetics. This leads to a cellular objective function

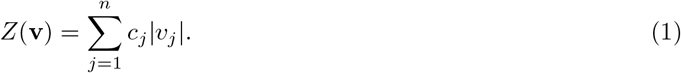

In the idealized case, **c** would be an indicator vector whose *j*-th entry is 1 if the cell is trying to maximize the reaction *R*_*j*_ and is zero otherwise. In practice, since the true cellular objective function is unknown, we will interpret *c*_*j*_ ∈ [0, 1] as the probability that the cell is trying to maximize reaction *j*. FBA thus identifies the vector **v** within the solution space that describes the mass balanced metabolic network state most capable of engaging the objective biological processes under the given conditions by optimizing

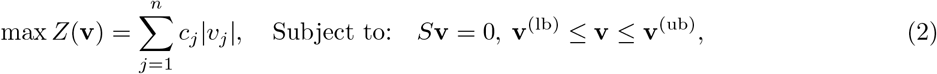

**v**^(lb)^ and **v**^(ub)^ are vectors with entries 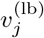 and 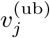 and the inequalities are defined componentwise.

Most FBA applications take a single network, where **c** and other parameters are known, and model phenotypic effects of modulating those parameters such as simulating a genetic/pharmacological perturbation or identifying optimal conditions to maximize some process (e.g., production of a specific molecule). FBA is most effective when reaction rates are predicted using a known objective function and constraints. Under well-studied conditions, we may safely assume certain biological objectives, such as a bacterium in nutrient-rich growth media optimizing growth and biomass accumulation. However, when environmental variables change or cells are acting cooperatively in a multicellular system, these objectives change and become difficult to determine a priori.

One possible approach might be to try to estimate **c** and the corresponding reaction flux rates **v** based on gene product levels. The relationship between gene product levels and reaction flux in a reaction *R*_*j*_ is described by the Michaelis-Menten equation,

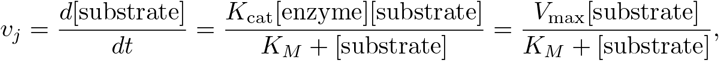

where the reaction rate *v*_*j*_ is a non-linear function of the substrate concentration [substrate], the concentration [enzyme] and experimentally-defined kinetic rate constant *K*_cat_ for the catalyzing enzyme, and the Michaelis constant *K*_*M*_ , the later two being commonly expressed using the product term *V*_max_. Enzyme concentration is subject to regulatory processes beyond transcription that include post-transcriptional/translational modification, stability, and turnover. Additionally, *K*_cat_ can only be derived experimentally and is not available for many enzymes, making it difficult to implement in large genome-scale metabolic network models. For these reasons, gene expression values are typically an unreliable predictor of reaction rates.

As opposed to traditional applications of FBA, we are less concerned with the precision of flux rate estimations and more interested in understanding the relative engagement of metabolic pathways by cells. We therefore consider dynamic gene expression changes between a given cell and a specified reference state, using relative expression values as a metric of active regulation in order to learn a cellular objective function and concordant network state (i.e., first learn **c** and then learn **v**). In summary, our goals are two-fold:

1. First, we aim to learn a vector **c** that describes which reactions the cell is trying to maximize.
2. Then, given **c**, we aim to find **v** by solving the constrained optimization problem (2).

## 3 Related Work

Historically, metabolic phenotypes in transcriptional data have been identified using differential gene expression with pathway enrichment methods such as GSEA [27]. Some methods are specifically designed to focus on metabolic pathways such as KEGG [11], MetaboAnalyst [17], or Reactome [4]. More recently, various methods have been developed to integrate transcriptomic data and constraint-based modeling. For bulk population analysis, variations on FBA such as parsimonious FBA (pFBA), dynamic FBA (dFBA), or flux variability analysis (FVA) are able to incorporate gene expression information, as reviewed in [24].

Compass [28], a recently developed method, provides an extension of gene expression-constrained FBA to single-cell transcriptomics. In order to circumvent the previously discussed caveats with defining cellular objective functions in mammalian cells, i.e., that **c** is unknown, Compass instead estimates a score for each reaction in each cell representing the propensity of a cell to engage that reaction using serial rounds of linear optimization. First, an initial set of transcription-agnostic ‘maximal fluxes’ 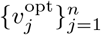 is calculated using network stoichiometries and user-defined parameters such as nutrient uptake limits. Gene expression values are then mapped to reactions and converted into penalty scores, where low expression of genes involved in a reaction incurs a high penalty, and, for every reaction *R*_*j*_ in every cell *i*, Compass solves a linear optimization problem that minimizes reaction penalties (i.e., minimizes use of lowly expressed genes) while preserving a specified proportion *ω* of the maximum flux, such that 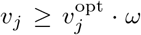. The magnitude of *v*_*j*_ represents the maximal flux that reaction *R*_*j*_ may carry with minimal resistance from the cell, where 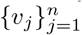 can be inverted and scaled within each cell to produce a set of scores signifying the relative capacity of a given cell to carry flux through each reaction.

While Compass does avoid the pitfalls of estimating metabolic state using a biologically irrelevant objective function such as biomass accumulation and does notably capture both known and novel biochemical phenotypes, iteration through each reaction in each cell generates a substantial computational cost and renders scalability difficult. Additionally, the capacity of a cell to engage a metabolic reaction transcriptionally does not necessarily imply a biological imperative to do so. For this reason, we hypothesize that identifying a cellular objective function based on dynamic gene expression will provide useful information about metabolic programs that mechanistically drive processes such as cellular differentiation and effector functionality, while simultaneously mitigating the computational burden of serial linear optimization.

To address these challenges, we propose a deep learning method for high-throughput prediction of metabolic network activity from scRNAseq data. We first build an (undirected) graph *G* = (*V, E*) of a given metabolic network where the nodes *V* represent reactions and edges *E* represent the production/consumption of a metabolite. Here, we define edges {*v*_*i*_, *v*_*j*_}∈ *E* if either reaction generates a product that the other consumes as a substrate, and edge weights represent metabolite flux, or the amount of metabolite moving between nodes *v*_*i*_ and *v*_*j*_ per arbitrary unit of time. We define node features as the relative gene expression between a given cell and a specified reference cell, which we map to the reactions catalyzed by each gene product to obtain a vertex-weighted graph that can be used to learn cellular objective functions and subsequent flux-balanced network state solutions.

## 4 GEFMAP: Gene Expression-based Flux Mapping and Metabolic Pathway Prediction

In this section, we outline our method, which is illustrated in Figure 1. Our approach is based on using a GNN to find a large, highly connected subgraph in a graph derived from the metabolic network, using the output of this GNN to derive a cellular objective function, and then solving this objective function via neural network which is geometrically constrained in order to preserve mass balance in the system.

**Fig. 1.**
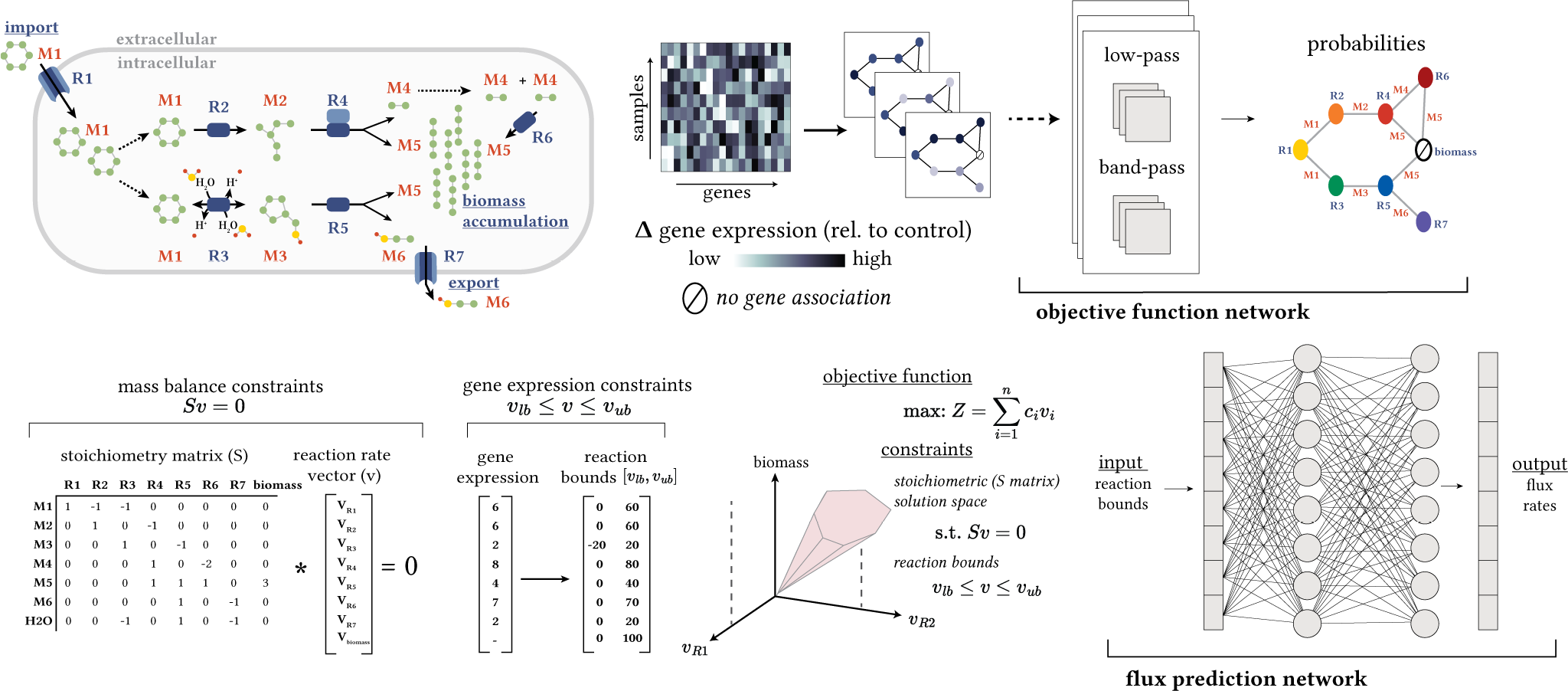
Our method is based upon (1) constructing a graph from the metabolic network, (2) determining which reactions a cell is trying to maximize via a graph neural network (3) using stoichiometric constraints to formulate an objective function and (4) using a neural network to find flux rates which optimize this objective.

### 4.1 Metabolic Network Graph Generation

Given a network with *m* metabolites and *n* reactions, we treat the nodes of the graph as the reactions and the edges of the graph as the amount of metabolites flowing through them. We let *Ŝ* denote a Boolean stoichiometry matrix defined by

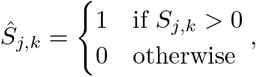

and we calculate an *n × n* reaction adjacency matrix, *A* defined by

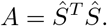

We note that *A*_*j*,*k*_ is nonzero if and only if reactions *R*_*j*_ and *R*_*k*_ utilize a common metabolite in a producer/consumer relationship. The resulting graph represents the connectivity of the metabolic network.

Nodes corresponding to reactions that are catalyzed by a gene-encoded enzyme have gene expression features, and we assign values *w*_*j*_ to each node using the reaction gene associations provided in the genomescale models (GEM). If a reaction can be catalyzed by multiple enzymes separately (A or B), we use the maximum expression value for the set of genes, and if the enzymes form a complex (A and B), we take the minimum expression value. We apply *ℓ*^1^-normalization and min-max scaling to reaction expression values; reactions without gene associations are assigned a value of 0 for estimating the objective function and 1 for predicting the flux rates.

### 4.2 Inferring the objective function

The first key component of GEFMAP is a graph neural network which uses the geometry of the metabolic network to infer the objective **c** in (2). Our approach is based on [18], which used a GNN to find the maximum clique (largest fully connected subgraph) in a given graph. Here, we consider the maximum weighted clique, which is defined as

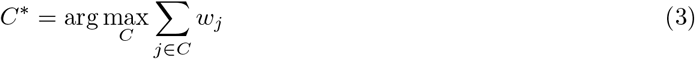

where the max is taken over all cliques, i.e., sets *C* ⊆ *V* such that *A*_*j*,*k*_ *>* 0 for all *j, k* ∈ *C*, and the node weights *w*_*j*_ are the normalized reaction expression features described in the previous subsection. The input to our GNN is an *n ×* 4 feature matrix *X* where the columns of *X* represent the eccentricity, clustering coefficient, node weight, and weighted degree of the nodes. (This differs slightly from [18] which used an *n ×* 3 matrix with unweighted node degree and omitted the node weight column.) The output of this GNN is vector **p** where *p*_*j*_ is to be thought of as the *probability* that the *j*-th node is a member of *C*^∗^.

In [18], after computing **p**, the authors apply a greedy decoder. This greedy decoder uses **p** to construct *K* different cliques *C*_1_, … *C*_*K*_ and then chooses *C*^∗^ to be largest *C*_*k*_. (In [18], largest would mean most vertices, whereas here it would mean largest value of 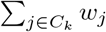). Each of the cliques *C*_*k*_ are formed in an iterative manner. We first order the nodes according to the probability that they are included in the max weighted clique *p*_*j*_. To construct the first clique *C*_1_, we pick a seed vertex *j* with the maximal value of *p*_*j*_ and initialize *C*_1_ = {*j*}. Then, at each step, we consider a node *j*^*′*^ such that 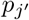 is as large as possible amongst all nodes which have not previously been considered. If *C*_1_ ∪{*j*^*′*^} forms a clique, we add *j*^*′*^ to *C*_1_. Otherwise, we discard *j*^*′*^ and move on to the next candidate and proceed in this manner until all nodes have been considered. Then, having constructed the first *k* cliques, 1 ≤*k* ≤*K* −1 , the *k* + 1 -st clique *C*_*k*+1_ i s constructed via a similar procedure as used to construct *C*_1_. The only differences are that (i) the seed node *j* is chosen to be as large as possible amongst all nodes which have not previously been used as a seed node and (ii) all nodes which have previously been used as the seed node are automatically excluded from the clique.

The GNN used to compute **p** is based on the method introduced in [29]. In each layer, the network uses two types of filters: (i) low-pass filters inspired by Kipf and Welling’s Graph Convolutional Network (GCN) [9] and (ii) band-pass wavelet filters inspired by the geometric scattering transform [6] (see also [5,32]). The low-pass filters, which are similar to those used in standard message-passing neural networks (see, e.g., [30]) are constructed by applying powers of the normalized adjacency matrix and aim to smooth the node features, i.e., to ensure that the GNN’s representation of node *j* is similar to node *k* if there is an edge between *j* and *k*. The band-pass filters, which are constructed by subtracting random-walk matrices raised to different powers, on the other hand, aim to prevent too much smoothing and capture important oscillations in the node features. To understand the importance of these band-pass filters and their capacity to capture different information than the GCN filters, consider the case where *C* ^∗^ consists of 30 nodes and some n ode *j* is a neighbor of 29 of them. A network which is solely based on low-pass smoothing operations will produce a hidden representation of node *j* which is similar to its neighbors. Therefore, it is highly likely to mistakenly think that *j* is a member of the clique. By contrast, the band-pass filters do not smooth the node features and allow the network to detect some manner in which *x*_*j*_ is different than the values of **x** at the neighbors of *j*.

Since the low-pass and band-pass filters can be seen as serving competing goals, promoting smoothness in features versus preserving informative oscillations, we use a localized attention mechanism to balance the importance of the various filters in the network. For each filter *f*, the attention mechanism computes a score vector ***α***_*f*_ (where (***α***_*f*_ )_*j*_ is the importance of filter *f* and node *j* ) and uses these scores to reweight the hidden representation of each vertex. The network then passes this hidden representation through a multi-layer perceptron (MLP) before applying another GNN layer. For further details on the layers utilized in this GNN (which also includes readout layers and an initial transformation of the node features parameterized by an MLP), we refer the readers to Section 3 of [18].

Unlike more common GNN tasks such as node-classification, there is no obvious loss function to use while training the GNN to solve (3). In [18], the authors used a two part loss function 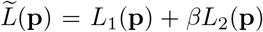 (where *β* is a tunable parameter). The first part of the loss function, which is defined

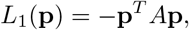

encourages the network to find a vector **p** whose mass is concentrated on connected nodes. The second term is given by

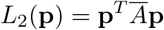

where 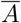 is the adjacency matrix of the complement graph, i.e., for *i ≠ j*, 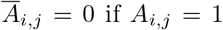 (and vice versa). Lemma 1 of [18] shows that *L*_2_(**p**) is equal to zero if and only if the support of **p** is contained in a clique. Therefore, minimizing both of these loss functions jointly ensures that we find a probability vector **p** whose support is concentrated in a clique while containing as many connections as possible. Here, we wish to ensure that the support of **p** is concentrated in a set *C*^∗^ such that 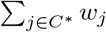 is as large as possible. Therefore, we modify the loss function used in [18] and utilize a new loss function defined by

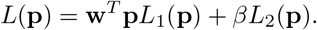

We note that in the idealized case where *p*_*j*_ = 1 for *j* ∈ *C*^∗^ and *p*_*j*_ = 0 otherwise, the term **w**^*T*^ **p** becomes the quantity that we are trying to optimize 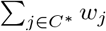. Therefore, this modification to the loss function leads the network to find a clique with high-levels of gene expression.

After finding the maximum clique *C*^∗^, a natural choice would be to let **c** be the corresponding indicator function, i.e., *c*_*i*_ = 1 if *i* ∈ *C*^∗^, *c*_*i*_ = 0 otherwise. However, we find that the stringency of the objective function resulting from this method is not highly reflective of the cell biology for a number of reasons. Metabolic networks are relatively sparse and often include reactions such as nutrient import or transport between eukaryotic organelles that may have only a few adjacent reactions but regulate highly-connected subnetworks. Additionally, such reactions are often transcriptionally regulated, for example, the solute-carrier (SLC) or ATP-binding-cassette (ABC) family of transporters, and are of particular interest because of their small-molecule drug targetability [23]. Therefore, we instead consider a relaxation of the maximum clique problem and simply set **c** = **p**. Indeed, the constraint that **c** is supported in a clique is overly rigid. While we do expect the reactions which a cell is trying to optimize to form a highly connected subnetwork, we do not expect them to form a clique per se. With this framework, we then roughly interpret *c*_*i*_ = *p*_*i*_ as the probability that the cell is trying to maximize reaction *i*.

### 4.3 Null Space Network for Solving the Objective Function

The second key component of GEFMAP is a neural network which maximizes Σ_*i*_ *c*_*i*_ |*v*_*i*_| over **v** lying in the null space of *S*. In order to account for the geometric constraint, *S***v** = 0, we first compute an orthonormal basis for Null(*S*), **b**_**1**_, … , **b**_**K**_, and note that any **v** ∈ Null(*S*) may be written as

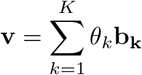

for some *θ*_1_, … , *θ*_*k*_ ∈ ℝ. Therefore, we train an MLP with three layers and ReLU activations to learn the *θ*_*k*_. We note that one of the objectives of this network is to bypass the use of a linear program for each cell, since each cell can have a unique objective. However, we do use a linear program in the process of generating our training data.

## 5 Experiments

### 5.1 Experimental Setup

Extensive experiments on GEFMAP are performed on a synthetic toy data set, an augmented E. coli data set, and a human embryoid body data set to validate the model’s efficacy in modeling complex metabolic networks. For code needed to reproduce our experiments, please see https://github.com/KrishnaswamyLab/metabolic_GNN. The first subnetwork of GEFMAP computes the cell’s metabolic objective and this is used by the second subnetwork to solve for it. All of the graphs generated from these data sets are treated as undirected by all the networks except in the case of one of the baselines MagNet [31], which is a directed graph neural network that is capable of dealing with directed edges (where the direction of the edge corresponds to the direction of the reaction). For all of the methods, we use the upper and lower bounds **v**^ub^ and **v**^lb^ as input features. As a loss function, we use the mean squared error between **v** and the optimal ground truth solution **v**^∗^ (where the ground truth vectors are generated via a direct linear solver).

#### Baselines

As baselines for comparison, we consider several other deep-learning architectures in place of the null space network. Our first baseline is the graph convolutional network (GCN) [9], a widely used message-passing network for node-level tasks on undirected graphs. As a second baseline, we also consider MagNet[31], which aims to incorporate directional information into the geometric deep learning framework. To apply MagNet, we view the metabolic network as a directed graph, where edge direction corresponds to the direction of the flow, i.e., there is a directed edge from *R*_*i*_ to *R*_*j*_ if *R*_*i*_ produces a metabolite which is then utilized by *R*_*j*_. Given the directed graph, MagNet then uses a complex Hermitian matrix known as the magnetic Laplacian[2,14,19] to define a notion of convolution on a directed graph (building off of several previous works, such as [3,12], which uses the standard graph Laplacian on an undirected graph). As a final baseline, we use a simple multilayer perceptron (MLP) that does not use the geometry of the metabolic network.

### 5.2 Validating our Objective Function on Core E. Coli Network

Here, we evaluate the biological relevance of the objective function introduced in Section 4.2, which is based on a relaxed formulation of the max-weighted clique problem, versus biomass accumulation, a generic objective function not based on biological parameters. We first test our methods using gene expression from E. coli cultured under various conditions [25]. The original data contained over 250 samples and approximately 150 experimental conditions, which we augmented to over 3000 samples by adding Gaussian noise to gene expression values. For these experiments, we use absolute rather than relative expression values due to the experimental absence of a reference transcriptome. Using the E. coli core metabolic model [21], we construct a *m × n* reaction adjacency matrix *A* representative of *m* = 72 metabolites and *n* = 95 reactions. Expression of 137 genes was mapped onto the 95 reactions, examples illustrated in (Fig. 2a), and reaction expression values were used to compute **c**.

**Fig. 2.**
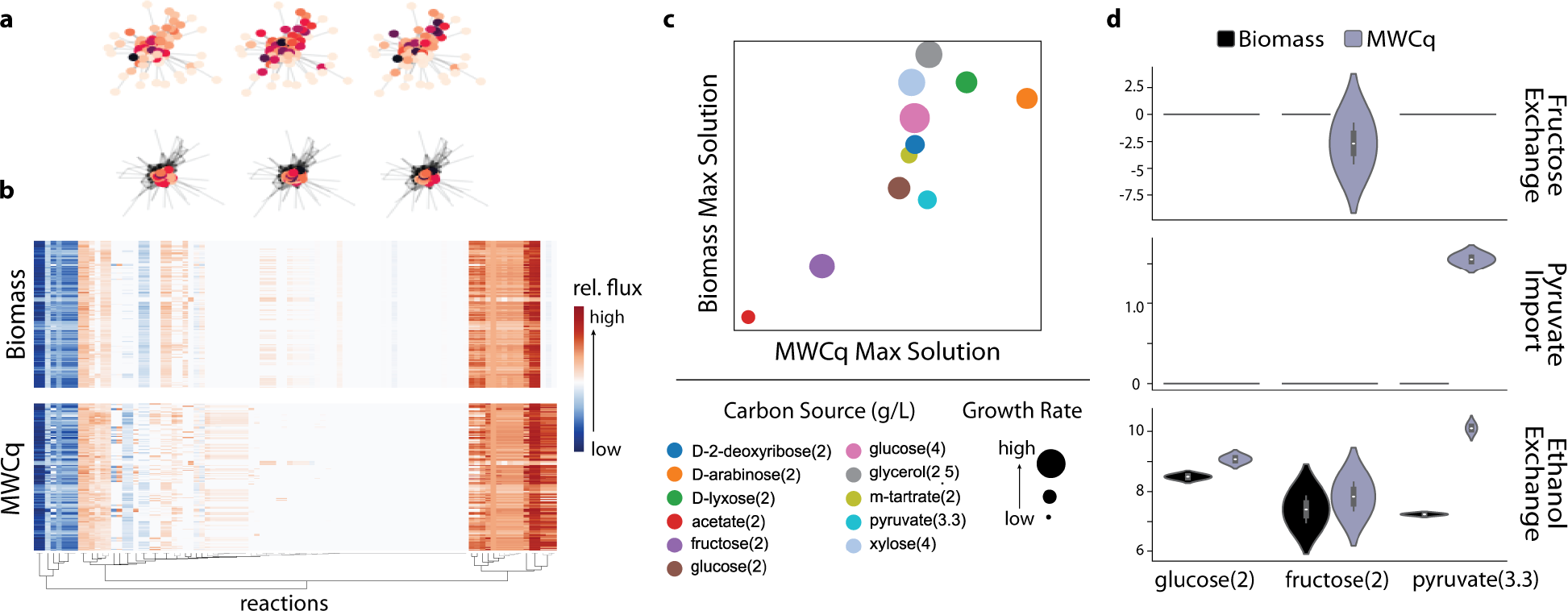
Max weighted clique (MWCq) based cellular objective function. (a) Illustration of vertex-weighted metabolic network graphs (left) constructed from the core E. coli metabolic network and associated max-weighted cliques (right). (b.) Predicted reaction rate values from E. coli RNA sequencing samples to maximize the flux through highly-connected subnetworks (top) or biomass/growth objectives (bottom). (c.) Correlation between maximal flux rates for different objectives. Datapoints are colored by experimental growth condition and datapoint size represents experimentally-measured relative growth rates. (d.) Predicted flux rates for reactions involved in central carbon metabolism for bacteria grown in the presence of either glucose, fructose, or pyruvate.

To test the result of the relaxed max-clique objective function (in isolation of the null space network), we next used FBA, with an off-the-shelf linear solver, to estimate reaction flux rates across the network that maximized either flux through the relaxed max clique or, as a comparison, maximized biomass accumulation (growth) (Fig. 2b). When maximizing with respect to our max-clique based objective, we observe more diversity across samples than when maximizing biomass. We hypothesize that this is representative of pathways engaged by cells not in states of heavy growth. Therefore, we subset samples by carbon source (Fig. 2c) and compare how experimentally-measured growth rate correlates with the predicted maximized flux rates when using relaxed max clique vs. biomass.

For both objective functions, we observed a general trend that bacteria with higher measured growth rates were predicted to have higher flux rates. However, when optimizing for biomass, there are a number of samples for which we are able to find transcriptionally-constrained flux solutions that predict high levels of biomass increase despite the fact that the measured growth rates are low, reinforcing the need for a more physiologically relevant objective function. To further explore this, we next look specifically at bacteria utilizing either 2 g/L glucose, fructose, or pyruvate and compare differences in the predicted rate of reactions involved in these different components of central carbon metabolism (Fig. 2d). For both fructose and pyruvate, the only conditions in which we observe predicted activity in the reactions that mediate cellular import are in the presence of the nutrient and with solutions that maximize flux through the relaxed max clique, suggesting that this method is able to capture the cellular metabolic response to environmental nutrient availability.

### 5.3 Learning FBA Solution Flux Estimations

For the purposes of validating our null space network, we generated a synthetic data set comprised of 892 graphs with each graph having *n* = 8 reactions and *m* = 12 metabolites. Gene expression values were chosen by random sampling of a normal distribution, and ground truth reaction flux rates were determined by flux balance analysis (FBA) using an arbitrary objective function. We additionally test the augmented E. coli data that consists of over 3000 graphs with 95 reactions, 72 metabolites, and 2204 edges where the ground truth reaction flux rates were computed using the geometric scattering GNN. We evaluate accuracy via the metric used to compare the Pearson Correlation Coefficient (PCC) between predicted flux values and the ground truth for both the null space network and the baseline models.

The results in Table 1 show the mean *±* standard deviation PCC of the models, run five times on each data set, employing different random states each time during the train-test split process. GCN performs considerably better than the simple MLP, due to its ability to incorporate the network structure of the data, and MagNet performs better than GCN likely due to its ability to incorporate the direction of the reactions. The null space network, which also incorporates the structure of the network via the constraint *S***v**= 0, is the top performing model on both data sets.

**Table 1.** Results of flux predictions (PCC with ground truth) over five runs (mean *±* standard deviation). The best-performing method is highlighted in bold.

| Data | Synthetic | *E. coli* |
| --- | --- | --- |
| Model name |  |  |
| Null Space | **$0.928 \pm 0.016$** | **$0.939 \pm 0.001$** |
| MagNet | $0.841 \pm 0.014$ | $0.920 \pm 0.005$ |
| GCN | $0.643 \pm 0.021$ | $0.741 \pm 0.019$ |
| MLP | $0.308 \pm 0.011$ | $0.409 \pm 0.004$ |

### 5.4 Human Embryoid Networks

To test the strength of GEFMAP on real-world data, we use a previously published scRNAseq data set of human embryonic stem cells (hESCs) gathered at 5 timepoints over 27 days [20]. We generate a human metabolic network graph using Recond3 [1] that consists of 5835 metabolites and 10600 reactions. We consider dynamic gene expression between a given timepoint and precursor embryonic stem (ES) cells sequenced at the initial timepoint (*t* = 0), which are used as the input for GEFMAP. We define an objective score as the likelihood values that each reaction is included in the max weighted clique, representing the relative likelihood a cell is actively engaging a given reaction.

We first aggregate reaction scores across all neuronal lineage cells and group reactions by pathways that were manually curated using Metabolic Atlas [13] (Fig. 3a). Of the pathways analyzed, we observe the highest objective scores in reactions involved in ketolysis (ketone consumption), as well as notable trends towards higher scores in TCA/OXPHOS (tricarboxylic acid cycle and oxidative phosphorylation) and REDOX (oxidation-reduction) regulation.

**Fig. 3.**
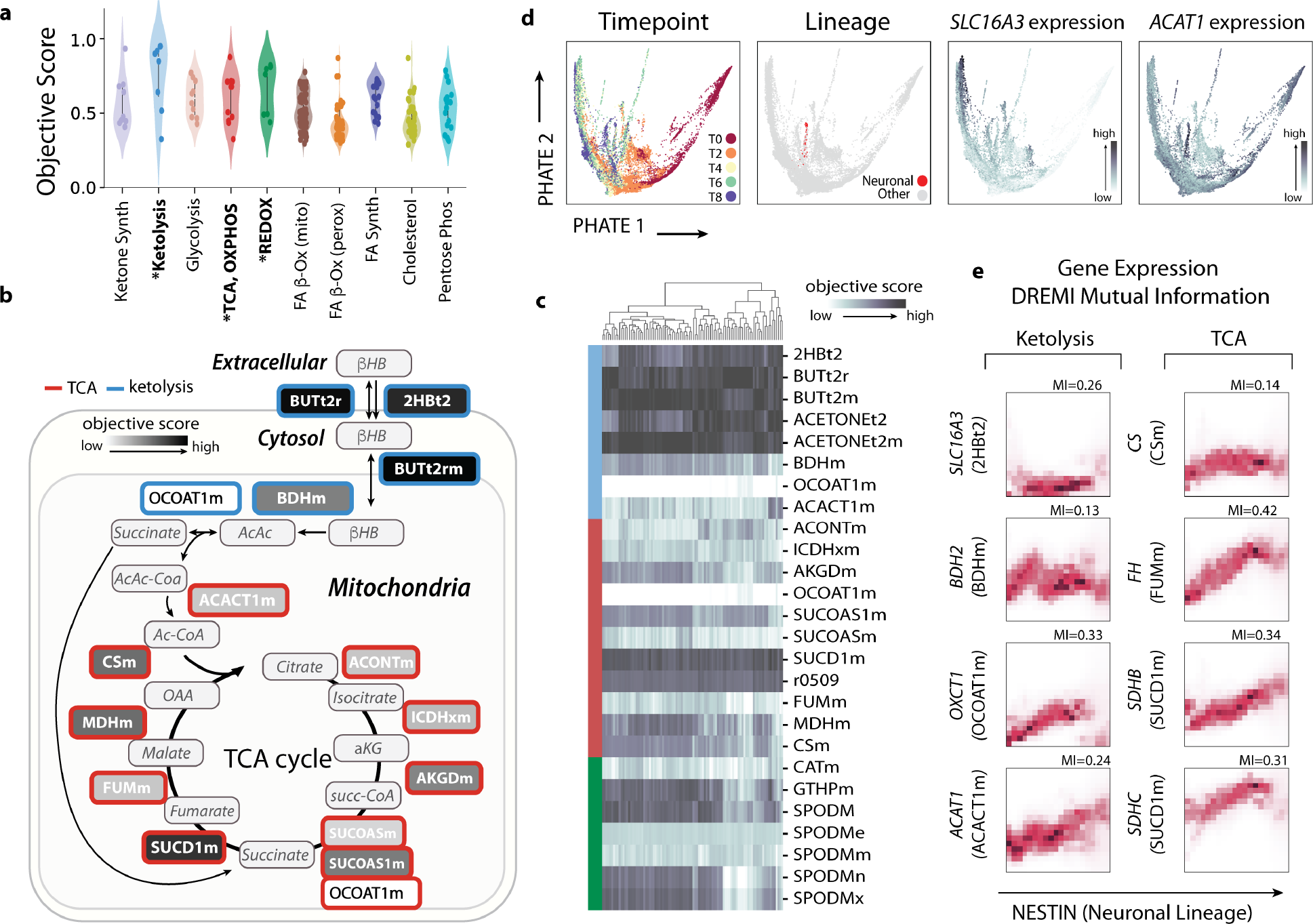
Human embryoid body data implicates ketone metabolism in regulation of neuronal lineage differentiation. (a.) Objective score (MWCq prediction) values for different metabolic pathways. Individual data points represent reactions, and objective score values represent median score across all cells. (b.) Illustration showing catabolism of the ketone *β*-hydroxybutrate (BHB) for utilization as a carbon substrate for the TCA. Reaction border color denotes pathway associate (blue = ketone catabolism, red = TCA) (c) Heatmap showing objective scores for individual reactions. Left side bar denotes pathway association (blue = ketone catabolism, red = TCA, green = REDOX). (d) PHATE embeddings [20] colored by experimental timepoint (far left), neuronal lineage identity (center left), and expression of ketone metabolic genes *SLC16A3* (*MPC4* ) (center right) and *ACAT1* (far right). (e) DREMI/DREVI [10] mutual information shows the relationship between expression of the neuronal lineage marker *NES* (*Nestin*) and expression of genes catalyzing reactions involved in ketolytis (left column) and the TCA (right column).

Ketone bodies (ketones) *β*-hydroxybutyrate (BHB) and acetoacetate (AcAc) are short-chain fatty acids that have recently gained public attention due to the growing popularity of ketogenic diets. These diets mimic natural fasting states, where ketones synthesized mainly in the liver provide an alternative fuel source to glucose [7]. Ketones are actively imported into the cell via monocarboxylate transporters (MCT) and transported into the mitochondria, where they may be converted to acetyl-CoA and enter the TCA for production of energy via ATP or anapleurotic intermediates (Fig. 3b). During fasting/starvation, ketone utilization is prioritized in tissues with high energy demands including brain, heart, and skeletal muscle, but has more recently been implicated in various other biological processes beyond energetics during both fasting and feeding in many diverse cell types [7,16,26].

Ketone catabolism is particularly intriguing in neuronal lineage cells. A well-documented phenotype of humans and animal models fed a ketogenic diet is general improvement in cognitive function as well as reduced pathology during neuronal damage or neurodegenerative disease [16]. Mechanistically, these phenotypes have been attributed to neuronal preference for BHB as a carbon source for mitochondrial metabolism, however alternative roles have been described that include regulation of signaling pathways and epigenetic modifications that enhance cellular function and stress resistance [16].

Neuronal differentiation has been shown to involve a concomitant increase in mitochondrial biogenesis and decrease in glycolysis and fatty acid oxidation, leaving ketones as a likely fuel source [15]. Indeed, we observe lower objective scores in glycolysis and fatty acid oxidation consistent with the literature (Fig. 3a). Within the ketolytic pathway, objective scores vary between individual reactions and across developing cells (Fig. 3c), suggesting specific reaction-level regulation during developmental processes. The highest scores are found in reactions that move BHB between cellular compartments (Fig. 3c).

To further explore how components of the ketolytic pathway are dynamically regulated during neuronal lineage development, we embed cells based on transcriptional similarity using PHATE [20] (Fig. 3d). We identify neuronal lineage cells using the lineage marker *ONECUT2* and visualize expression of the monocarboxylate transporter *SLC16A3* (*MCT4* ) that imports BHB into the cell (reaction 2HBt2) and the enzyme *ACAT1* that reversibly converts acetoacetyl-Coa into 2 acetyl-CoA (reaction ACACT1m) (Fig. 3d). We observe an overall trend in upregulation of *SLC16A3* at later experimental timepoints in the EB data , while *ACAT1* tends to decrease over experimental time with the exception of the neuronal lineage population. To more directly address the dynamics of ketyolytic gene expression within neuronal development, we use DREMI/DREVI [10] to estimate and visualize mutual information shared between these genes and the neural lineage marker *NES* (*Nestin*)(Fig. 3e). There is an generally positive correlation between *NES* expression and ketyolytic gene expression while patterns vary. For example *SLC16A3* expression is only high in *NES* ^high^ cells, while *ACAT1* continuously increases with *NES* expression. *BDH2* and *OXCT1* are most higly expressed in *NES* ^mid^ then begin to decrease again, which is the pattern that we see in most TCA genes.

In the mitochondria, BHB-derived carbon can enter the TCA as either succinate or Acetyl-CoA (Ac-CoA), however our model predicts a relatively low likelihood score for the reaction OCOAT1m that commits BHB-derived carbon to the TCA. Of note, we do not interpret this to mean that this reaction is not active in the cell, but rather that it is not transcriptionally engaged for developmental progression. Within the TCA, the reaction that we observe the highest likelihood score in is SUCD1m, which converts succinate to fumarate for the dual purpose for moving the TCA forward as well and simultaneously supplying electrons to the electron transport chain (ETC) via complex II. During development, the energetic requirements of proliferation and differentiation impose a heavy workload on the mitochondria, which may lead cells to increase activity of reactions such as SUCD1m to produce more ATP. However, rapid transport of electrons can lead to production of reaction oxygen species (ROS), which can damage DNA, membranes, and cause significant cell stress. Indeed, ketolysis has been implicated in protection from oxidative stress in part through activation of *NRF2* [15,7]. Because we observed high objective scores for reactions involved in REDOX regulation (Fig. 3a), an appealing hypothesis is that ketone metabolism supports neuronal differentiation energetically by increasing electron transport through complex II of the ETC and simultaneously protects cells from consequential oxidative stress either via *NRF2* or an alternative mechanism. In total, these results show that GEFMAP is capable of recapitulating known metabolic signatures as well as predicting novel reaction-level metabolic features from single cell transcriptomic data in human cells.

## 6 Conclusions and Future Work

In this paper, we infer metabolic states using geometric deep learning by introducing a novel method GEFMAP -Gene Expression-based Flux Mapping and Metabolic Pathway Prediction. GEFMAP is a two-part neural network where the first sub-network is a geometric scattering-based GNN that finds a large highly connected subnetwork with high levels of gene expression in order to determine the cell’s metabolic objective from upregulated sections of the transcriptomic profile and the second sub-network is a fully connected MLP that operates in the null space of the metabolic stoichiometric matrix to solve the objective to estimate the resultant flux rate. Experiments using on synthetic and complex real-world data sets show the ability of GEFMAP to determine the metabolic objective and estimate the flux rate with high accuracy. As a future direction, we will try to model the interplay between metabolomics and transcriptomics bidirectionally using metabolic measurements as well. Additionally, experimenting on human disease data in order to identify certain metabolic pathways that could impact treatment response is also a possible future direction to explore.

## 7 Acknowledgements

Yixuan He is supported by a Clarendon scholarship from the University of Oxford. Michael Perlmutter and Smita Krishnaswamy were partially funded by NSF DMS 2327211. Additionally, Smita Krishnaswamy was partially supported by NSF Career Grant 2047856, by NIH 1R01GM130847-01A1, and by NIH 1R01GM135929-01.

## Notes

https://github.com/KrishnaswamyLab/metabolic_GNN

### Competing Interest Statement

The authors have declared no competing interest.

## References

1. Brunk, E., Sahoo, S., Zielinski, D.C., Altunkaya, A., ger, A., Mih, N., Gatto, F., Nilsson, A., Preciat Gonzalez, G.A., Aurich, M.K. ć A., Sastry, A., Danielsdottir, A.D., Heinken, A., Noronha, A., Rose, P.W., Burley, S.K., Fleming, R.M.T., Nielsen, J., Thiele, I., Palsson, B.O .: Recon3D enables a three-dimensional view of gene variation in human metabolism. Nat Biotechnol 36(3), 272–281 (Mar 2018)

2. Cucuringu, M., Li, H., Sun, H., Zanetti, L.: Hermitian matrices for clustering directed graphs: insights and applications. In: International Conference on Artificial Intelligence and Statistics. pp. 983–992. PMLR (2020)

3. Defferrard, M., Bresson, X., Vandergheynst, P.: Convolutional neural networks on graphs with fast localized spectral filtering. In: Advances in Neural Information Processing Systems 29. pp. 3844–3852 (2016)

4. Fabregat, A., Jupe, S., Matthews, L., Sidiropoulos, K., Gillespie, M., Garapati, P., Haw, R., Jassal, B., Korninger, F., May, B., Milacic, M., Roca, C.D., Rothfels, K., Sevilla, C., Shamovsky, V., Shorser, S., Varusai, T., Viteri, G., Weiser, J., Wu, G., Stein, L., Hermjakob, H., D’Eustachio, P.: The reactome pathway knowledgebase. Nucleic Acids Research 46(D1), D649–D655 (Nov 2017). https://doi.org/10.1093/nar/gkx1132, 10.1093/nar/gkx1132

5. Gama, F., Ribeiro, A., Bruna, J.: Diffusion scattering transforms on graphs. In: 7th International Conference on Learning Representations, ICLR 2019 (2019)

6. Gao, F., Wolf, G., Hirn, M.: Geometric scattering for graph data analysis. In: International Conference on Machine Learning. pp. 2122–2131. PMLR (2019)

7. García-Rodríguez, D., Giménez-Cassina, A.: Ketone bodies in the brain beyond fuel metabolism: From excitability to gene expression and cell signaling. Frontiers in Molecular Neuroscience 14 (Aug 2021). https://doi.org/10.3389/fnmol.2021.732120, 10.3389/fnmol.2021.732120

8. Heirendt, L., Arreckx, S., Pfau, T., Mendoza, S.N., Richelle, A., Heinken, A., Haraldsdóttir, H.S., Wachowiak, J., Keating, S.M., Vlasov, V., Magnusdóttir, S., Ng, C.Y., Preciat, G., Žagare, A., Chan, S.H.J., Aurich, M.K., Clancy, C.M., Modamio, J., Sauls, J.T., Noronha, A., Bordbar, A., Cousins, B., Assal, D.C.E., Valcarcel, L.V., Apaolaza, I., Ghaderi, S., Ahookhosh, M., Guebila, M.B., Kostromins, A., Sompairac, N., Le, H.M., Ma, D., Sun, Y., Wang, L., Yurkovich, J.T., Oliveira, M.A.P., Vuong, P.T., Assal, L.P.E., Kuperstein, I., Zinovyev, A., Hinton, H.S., Bryant, W.A., Artacho, F.J.A., Planes, F.J., Stalidzans, E., Maass, A., Vempala, S., Hucka, M., Saunders, M.A., Maranas, C.D., Lewis, N.E., Sauter, T., Palsson, B.Ø., Thiele, I., Fleming, R.M.T.: Creation and analysis of biochemical constraint-based models using the COBRA toolbox v.3.0. Nature Protocols 14(3), 639–702 (Feb 2019). 10.1038/s41596-018-0098-2

9. Kipf, T.N., Welling, M.: Semi-supervised classification with graph convolutional networks. In: 5th International Conference on Learning Representations, ICLR 2017, Toulon, France, April 24-26, 2017, Conference Track Proceedings (2017)

10. Krishnaswamy, S., Spitzer, M.H., Mingueneau, M., Bendall, S.C., Litvin, O., Stone, E., Pe’er, D., Nolan, G.P.: Conditional density-based analysis of t cell signaling in single-cell data. Science 346(6213) (Nov 2014). https://doi.org/10.1126/science.1250689, 10.1126/science.1250689

11. Kyoto Encyclopedia of Genes and Genomes (KEGG): Kyoto encyclopedia of genes and genomes (Year), https://www.genome.jp/kegg/

12. Levie, R., Monti, F., Bresson, X., Bronstein, M.M.: Cayleynets: Graph convolutional neural networks with complex rational spectral filters. IEEE Trans. Signal Process. 67(1), 97–109 (2019). 10.1109/TSP.2018.2879624

13. Li, F., Chen, Y., Anton, M., Nielsen, J.: Gotenzymes: an extensive database of enzyme parameter predictions. Nucleic Acids Research 51(D1), D583–D586 (Sep 2022). https://doi.org/10.1093/nar/gkac831, 10.1093/nar/gkac831

14. Lieb, E.H., Loss, M.: Fluxes, Laplacians, and Kasteleyn’s theorem. In: Statistical Mechanics, pp. 457–483. Springer (1993)

15. Maffezzini, C., Calvo-Garrido, J., Wredenberg, A., Freyer, C.: Metabolic regulation of neurodifferentiation in the adult brain. Cellular and Molecular Life Sciences 77(13), 2483–2496 (Jan 2020). https://doi.org/10.1007/s00018-019-03430-9, 10.1007/s00018-019-03430-9

16. Mattson, M.P., Moehl, K., Ghena, N., Schmaedick, M., Cheng, A.: Intermittent metabolic switching, neuroplasticity and brain health. Nature Reviews Neuroscience 19(2), 81–94 (Jan 2018). https://doi.org/10.1038/nrn.2017.156, 10.1038/nrn.2017.156

17. MetaboAnalyst: Metaboanalyst: A comprehensive tool suite for metabolomic data analysis (Year), https://www.metaboanalyst.ca/

18. Min, Y., Wenkel, F., Perlmutter, M., Wolf, G.: Can hybrid geometric scattering networks help solve the maximum clique problem? In: NeurIPS (2022), http://papers.nips.cc/paper_files/paper/2022/hash/8ec88961d36d9a87ac24baf45402744f-Abstract-Conference.html

19. Mohar, B.: A new kind of Hermitian matrices for digraphs. Linear Algebra and its Applications 584, 343–352 (2020)

20. Moon, K.R., van Dijk, D., Wang, Z., Gigante, S., Burkhardt, D.B., Chen, W.S., Yim, K., van den Elzen, A., Hirn, M.J., Coifman, R.R., Ivanova, N.B., Wolf, G., Krishnaswamy, S.: Visualizing structure and transitions in high-dimensional biological data. Nature Biotechnology 37(12), 1482–1492 (Dec 2019). https://doi.org/10.1038/s41587-019-0336-3, 10.1038/s41587-019-0336-3

21. Orth, J.D., Fleming, R.M.T., Palsson, B.Ø.: Reconstruction and use of microbial metabolic networks: the core Escherichia coli metabolic model as an educational guide. EcoSal Plus 4(1) (Jan 2010). 10.1128/ecosalplus.10.2.1

22. Orth, J.D., Thiele, I., Palsson, B.Ø.: What is flux balance analysis? Nature Biotechnology 28(3), 245–248 (Mar 2010). 10.1038/nbt.1614

23. Rives, M.L., Javitch, J.A., Wickenden, A.D.: Potentiating SLC transporter activity: Emerging drug discovery opportunities. Biochemical Pharmacology 135, 1–11 (Jul 2017). https://doi.org/10.1016/j.bcp.2017.02.010, 10.1016/j.bcp.2017.02.010

24. Sahu, A., Blätke, M.A., Szymański, J.J., Töpfer, N.: Advances in flux balance analysis by integrating machine learning and mechanism-based models. Computational and Structural Biotechnology Journal 19, 4626–4640 (2021). 10.1016/j.csbj.2021.08.004

25. Sastry, A.V., Gao, Y., Szubin, R., Hefner, Y., Xu, S., Kim, D., Choudhary, K.S., Yang, L., King, Z.A., Palsson, B.O.: The Escherichia coli transcriptome mostly consists of independently regulated modules. Nature Communications 10(1) (Dec.). 10.1038/s41467-019-13483-w

26. Stubbs, B.J., Koutnik, A.P., Goldberg, E.L., Upadhyay, V., Turnbaugh, P.J., Verdin, E., Newman, J.C.: Investigating ketone bodies as immunometabolic countermeasures against respiratory viral infections. Med 1(1), 43–65 (Dec 2020). https://doi.org/10.1016/j.medj.2020.06.008, 10.1016/j.medj.2020.008.

27. Subramanian, A., Tamayo, P., Mootha, V.K., Mukherjee, S., Ebert, B.L., Gillette, M.A., Paulovich, A., Pomeroy, S.L., Golub, T.R., Lander, E.S., Mesirov, J.P.: Gene set enrichment analysis: A knowledge-based approach for interpreting genome-wide expression profiles. Proceedings of the National Academy of Sciences 102(43), 15545–15550 (2005). 10.1073/pnas.0506580102

28. Wagner, A., Wang, C., Fessler, J., DeTomaso, D., Avila-Pacheco, J., Kaminski, J., Zaghouani, S., Christian, E., Thakore, P., Schellhaass, B., Akama-Garren, E., Pierce, K., Singh, V., Ron-Harel, N., Douglas, V.P., Bod, L., Schnell, A., Puleston, D., Sobel, R.A., Haigis, M., Pearce, E.L., Soleimani, M., Clish, C., Regev, A., Kuchroo, V.K., Yosef, N.: Metabolic modeling of single th17 cells reveals regulators of autoimmunity. Cell 184(16), 4168–4185.e21 (Aug 2021). 10.1016/j.cell.2021.05.045

29. Wenkel, F., Min, Y., Hirn, M., Perlmutter, M., Wolf, G.: Overcoming oversmoothness in graph convolutional networks via hybrid scattering networks. arXiv:2201.08932 (2022)

30. Xu, K., Hu, W., Leskovec, J., Jegelka, S.: How powerful are graph neural networks? In: 7th International Conference on Learning Representations, ICLR 2019, New Orleans, LA, USA, May 6-9, 2019 (2019)

31. Zhang, X., He, Y., Brugnone, N., Perlmutter, M., Hirn, M.: Magnet: A neural network for directed graphs. Advances in neural information processing systems 34, 27003–27015 (2021)

32. Zou, D., Lerman, G.: Graph convolutional neural networks via scattering. Applied and Computational Harmonic Analysis 49(3), 1046–1074 (2020). 10.1016/j.acha.2019.06.003

